# Variance lower bound on fluorescence microscopy image denoising

**DOI:** 10.1101/2020.05.13.094748

**Authors:** Yilun Li, Sheng Liu, Fang Huang

**Affiliations:** Weldon School of Biomedical Engineering, Purdue University, West Lafayette, IN, USA; Purdue Institute for Integrative Neuroscience, Purdue University, West Lafayette, IN, USA; Purdue Institute of Inflammation, Immunology and Infectious Disease, Purdue University, West Lafayette, IN, USA

## Abstract

The signal to noise ratio of high-speed fluorescence microscopy is heavily influenced by photon counting noise and sensor noise due to the expected low photon budget. Denoising algorithms are developed to decrease these noise fluctuations in the microscopy data. One question arises: whether there exists a theoretical precision limit for the performance of a denoising algorithm. In this paper, combining Cramér-Rao Lower Bound with constraints and the low-pass-filter property of microscope systems, we develop a method providing a theoretical variance lower bound of microscopy image denoising. We show that this lower bound is influenced by photon count, readout noise, detection wavelength, effective pixel size and the numerical aperture of the microscope system. We demonstrate our development by comparing multiple state-of-the-art denoising algorithms to this bound. This theoretical bound provides a reference benchmark for microscopy denoising algorithms, and establishes a framework to incorporate additional prior knowledge into theoretical denoising performance limit calculation.

---

Rapid development of fluorescent microscopy techniques together with the availability of fast and sensitive cameras are enabling molecular observations at unprecedented spatial and temporal resolution. High-speed imaging of fluorescently tagged molecules suffers from low signal to noise ratio (SNR) due to the limited photon budget per fluorescent emitter. When ignoring sensor noise, the detected photons in each pixel follows Poisson distribution [1]. As a result, the SNR of a microscopy image decreases rapidly with decreasing number of detected photon. At these low light conditions, quantitative biological measurements result in imprecisions — fluctuations of measured signal mainly come from the uncertainty of photon detection rather than the underlying biological processes.

To improve the measurement precision, denoising algorithms are developed to decrease the noise fluctuation in the microscopy data caused by the photon detection process and the sensor. The key to such noise reducing capacity is the incorporation of additional prior knowledge or assumptions about the imaging system or biological specimens [2–7], for example, through exploiting the low pass filter property of a well-designed microscope [3] or assuming self-similarities of local structures in the specimens [4,7].

An ideal denoising algorithm will maintain the quantitative nature of microscopy images by providing estimations of pixel intensities with low variance and minimum biases or artifacts. However, common among the denoising algorithms that we tested, there exist tradeoffs between the estimation variance and the introduced bias. Typically, one could shift denoising performance with increased bias in trade for decreasing the denoising variance. Regarding this tradeoff, one fundamental question arises: How precise is precise enough? Is there a precision limit which a denoising algorithm could achieve at its best given a certain bias level? In this context, focusing on the case of unbiased denoising estimation of microscopy images, we derive an analytical expression to calculate the theoretical variance lower bound. This lower bound of variance provides a reference benchmark for the denoising algorithms and shows how the detected photons (e.g. their emission wavelength (*λ*) and expected photon counts) and the microscope system (e.g. numerical aperture (*NA*) of the objective lens, and pixel size of the camera) influence the precision limit of an unbiased microscopy denoising algorithm.

Our target here is to develop a variance lower bound for microscopy denoising algorithms by considering the common property of a far-field optical microscope system: the frequency response of a microscope is characterized by its optical transfer function (OTF) [8]. In a typical microscope system, the OTF boundary (with the radius of 2*NA*/*λ*) defines the region of detectable spatial frequencies whereas frequencies outside this boundary cannot transmit through the microscope system. Mathematically, this low-pass-filter property of microscope systems forms a rather strict constraint: The 2D Fourier transform of the underlying transmitted signal through a microscope must be zero outside the OTF boundary (Fig. 1). We expressed this constraint using the discrete Fourier transform matrix to form a set of equations corresponding to the vanishing components outside the OTF boundary in the following equation,

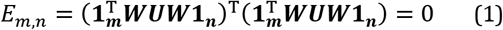

where (*m, n*) is the pixel coordinate outside the OTF boundary in the spatial frequency domain of a microscopy image (Fig. 1). Here *E_m,n_* is the magnitude square of the Fourier transform of the 2D image at (*m, n*), and superscript T denotes the conjugate transpose of a matrix. **1_m_** is a column vector with *m^th^* element equal to 1 while the others equal to 0, ***W*** is the discrete Fourier transform matrix [9] and ***U*** is the 2D array of the ideal image where each element *U_i,j_* is the expected photon counts of the pixel at position (*i, j*). Specifically, ***WUW*** is the 2D Fourier transform of an ideal image.

**Fig. 1.**
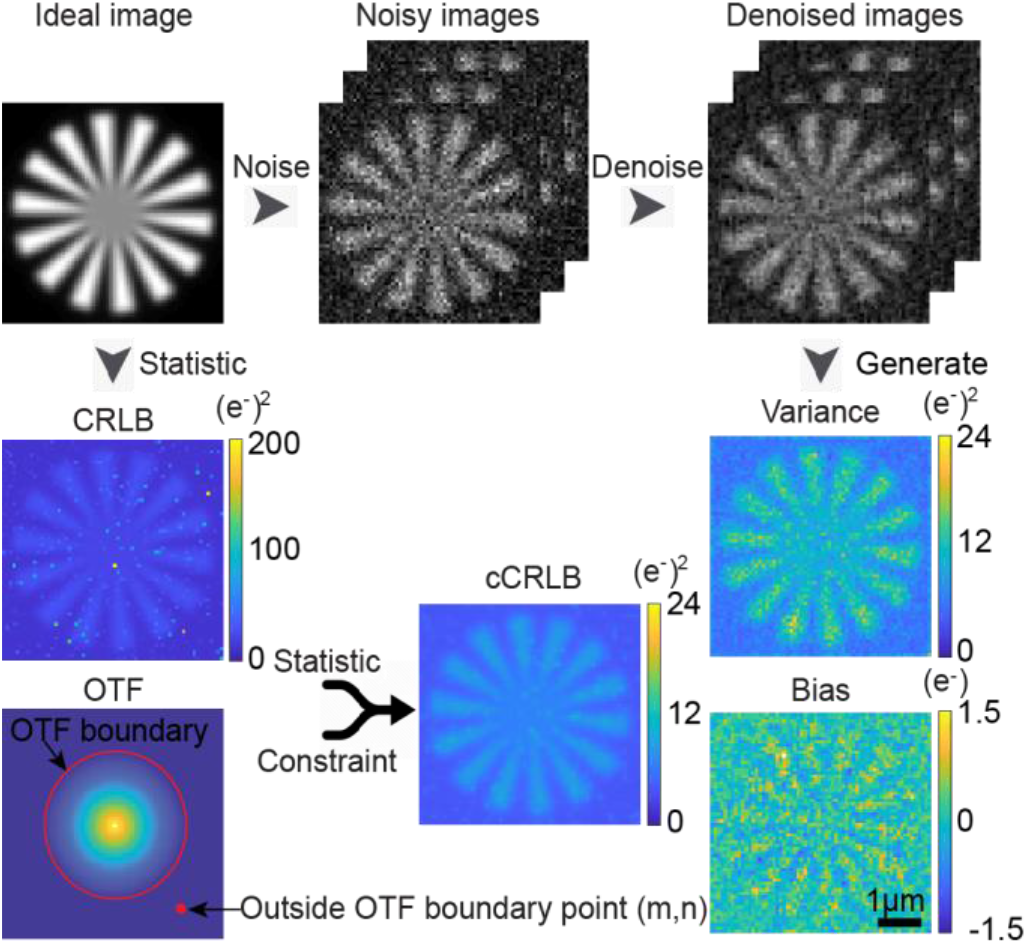
Schematic diagram of calculating cCRLB, bias, and variance of microscopy image denoising. CRLB is the variance lower bound on estimation of ideal image calculated from the noise model (Eq. 4–11). cCRLB is the variance lower bound on estimation of ideal image while taking CRLB and additional prior knowledge on frequency constraint (Eq. 1) into consideration. Bias map and variance map (on the right) were calculated pixel by pixel from 100 frames of denoised images.

To simplify notation, throughout this paper we used bold upper case letter to denote matrix (e.g. ***A***), bold lower case letter to denote vector (e.g. ***a***)and normal letter despite of upper or lower case to denote scalar (e.g. *a* or *A*). Moreover, all variables were denoted as letters in Italic while operations were denoted as normal letters (e.g. **A**).

Image denoising can be regarded as an estimation process during which one seek to estimate the noise-free ideal image from a detected noisy image with the help of additional knowledge. We considered the expected photon counts at different pixels as parameters to be estimated,

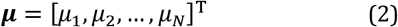

where ***μ*** = vec(***U***) is the vectorization form of ideal image ***U*** and *N* is the total number of pixels within a 2D image. Here we also defined the observation of the image,

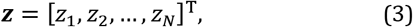

where *z_i_* is the observed readout of the *i^th^* pixel in the original noisy image.

To consider photon-counting noise and sensor noise, we assumed noise from each pixel is independent and can be modeled as the combination of two noise types [10]:

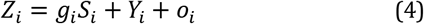

where *Z_i_, S_i_* and *Y_i_* are random variables each representing *i^th^* pixel’s readout, photon count and readout noise. *g_i_* stands for the pixel-dependent gain and *o_i_* is a constant offset pre-engineered into the readout process in order to prevent negative counts from camera sensors. In this paper, we assumed *S_i_* was a Poisson random variable with expectation *μ_i_* and *Y_i_* was a zero-mean Gaussian random variable with variance of 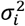,

Therefore, the probability of obtaining a specific camera readout *z_i_* (unit: analog to digit unit (ADU)) given an expected photon counts *μ_i_* (unit: e^-^) can be expressed as [10,11],

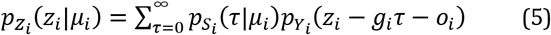

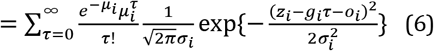

where *p_z_i__*() is the probability density function (PDF) of *Z_i_, p_S_i__*() is the probability mass function (PMF) of *S_i_* and *p_Y_i__*() is the PDF of *Y_i_*.

In estimation theory, Cramér-Rao Lower Bound [12] (CRLB) characterizes the lower bound on variance of unbiased estimators for parameters based on the likelihood function. In presence of constraints imposed on the parameters to be estimated, the lower bound on their variance can be further reduced since these constraints effectively reduce the dimensions of the parameter space [13]. This lower bound considering constraints on parameters is referred as constrained Cramér-Rao Lower Bound (cCRLB).

We defined the covariance matrix of the parameter vector,

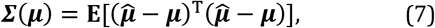

where **E** is taking the expectation over the distribution of observations ***z***.

According to Cramér and Rao et al [12,14,15]. the covariance matrix of the estimator satisfies the following inequation,

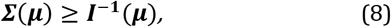

where matrix inequality means that ***∑***(***μ***) − ***I***^−1^(***μ***) is positive semidefinite and Fisher information matrix ***I***(***μ***) is an *N*×*N* matrix defined as,

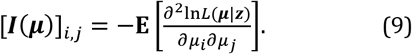

where *L*(***μ|z***) is the likelihood function of the image,

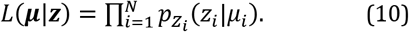

In our case, as we assumed the noise statistics at different pixels were independent, the off-diagonal elements of ***I***(***μ***) were zero while diagonal elements of ***I***(***μ***) can be calculated as [11],

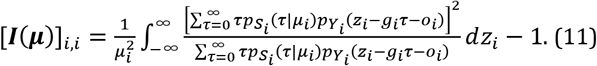

When we took the constraints into account, the constraint-lower bound (cCRLB) can be calculated as [13],

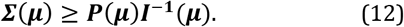

***P***(***μ***) is the projection matrix (shown in Ref. [13]) which takes the following form,

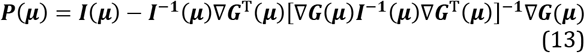

where ***G***(***μ***) is a column set of constraint functions satisfying,

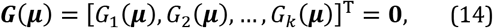

where *G_i_*(***μ***) is a scalar function of ***μ*** and *k* is the number of constraints. **0** refers to a column vector of all 0’s of size *k*.

Here, we adopt the convention that

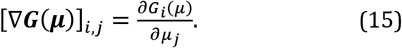

Next, we derived **∇*G***(***μ***) given our optical transfer function constraints in microscopy images as shown in Eq. 1. Combing Eqs. 1, 2 and 15 while changing the order of vectorization and derivation, each row of **∇*G***(***μ***) was calculated as,

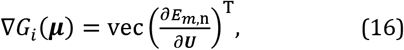

where pixel(*m, n*) in frequency domain is the *i^th^* pixel when we arranged all pixels outside OTF boundary in a raster scan order. We adopt the convention that

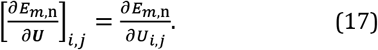

To calculate the above derivative matrix, we expanded *E_m,n_* from Eq. 1 as,

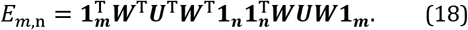

By defining

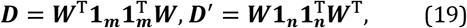

and using matrix derivative identities [16], we had

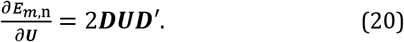

In summary, we calculated the unconstrained CRLB, ***I***^−1^(***μ***) (Eqs. 8 and 11), and then calculated the projection matrix, ***P***(***μ***) (Eqs. 13, 16 and 20). Finally, the cCRLB can be obtained by multiplying the resulting projection matrix with the unconstrained CRLB (Eq. 12). A schematic of our cCRLB calculation is shown in Fig. 1.

Here, we compared cCRLB with the achieved performance of three denoising algorithms: noise correction for scientific Complementary metal-oxide-semiconductor (sCMOS) camera (NCS) [3], non-local means filtering (NLM) [2] and automatic correction of sCMOS-related noise (ACsN) [4]. These algorithms rely on different principles: noise separations based on optical transfer function (OTF), self-similarity of local sub-regions within the image, and the combination of these two. To visualize and quantify the effectiveness of tested denoising algorithms, we evaluated their performances at a low signal-to-noise ratio (SNR) condition with peak expected photon count of 20 per pixel and a uniform background of expected photon count of 10 per pixel. We quantified denoising performance through calculating the achieved estimation variance as intensity fluctuation of the denoised image sequence and bias as the deviation between the ideal image and the mean of the denoised sequence. We showed that all three tested algorithms reduced the estimated variance substantially (variance reduction of 7.16-16.27 (e^-^)^2^ on average) compared to original noisy image with variance of 18.50 (e^-^)^2^ on average (Fig. 2). When examining both bias and variance of the denoising algorithms, we found NLM resulted in largest variance improvement (Fig. 2) achieving a denoised images’ variance of 2.23 (e^-^)^2^ on average but at the cost of significant higher absolute biases (~1.98 e^-^ on average at structure region defined as the region above average photon counts), causing the denoised image being statistically distorted from the ideal image. ACsN significantly reduced bias while achieving an estimation variance of 11.34 (e^-^)^2^ on average. NCS denoise algorithm achieved similar bias level as ACsN algorithm while achieving estimation variance of 8.87 (e^-^)^2^ on average. To demonstrate the precision improvement using denoising algorithms relative to the cCRLB, we calculated the pointwise ratio between achieved variance of a certain denoising algorithm and cCRLB. We found the resulting variance/cCRLB ratios of NCS were 1.56 on average (Fig. 2). The ratios of NLM were roughly 0.17 at the background regions while being 0.54 on average at the structure regions. AcSN had the ratio of 1.63 on average at the background region while being 2.24 on average at regions with structures.

**Fig. 2.**
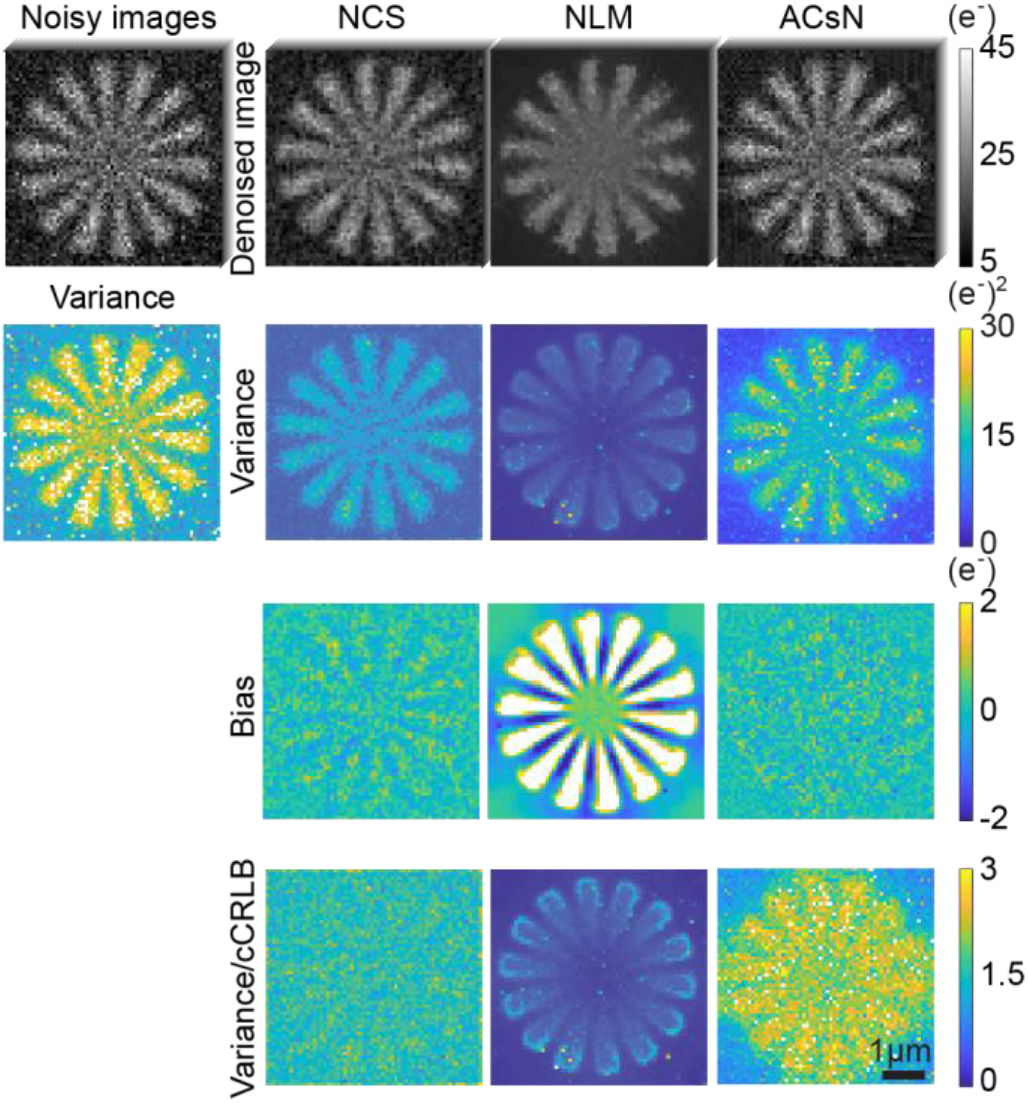
Comparison of three denoising algorithms. First row shows examples of noisy image (raw image before denoising) and denoised images from NCS, NLM and ACsN algorithms. Variance map (second row) and bias map (third row) were calculated using 100 frames of corresponding noisy or denoised images. Variance/cCRLB ratio maps (fourth row) were calculated from variance maps divided by cCRLB pixel by pixel.

The bias and variance/cCRLB ratios results were demonstrated in Fig. 3 where we grouped the bias and ratios based on pixels’ photon counts. NCS resulted in consistent performance across the tested image despite the variations of the expected photon counts. NLM resulted in increased bias and variance/cCRLB ratio in the positions with high photon counts than that in the low photon counts positions. ACsN maintained unbiased performance despite the variation of photon counts while increasing variance/cCRLB ratio with increased photon counts (Fig. 3b).

**Fig. 3.**
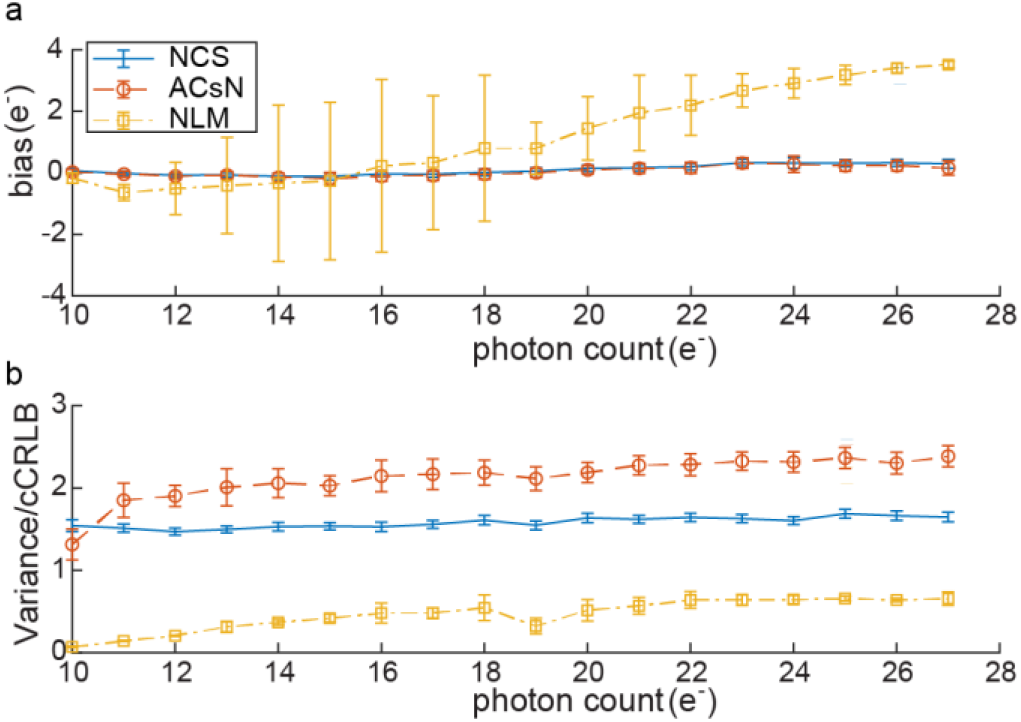
Comparison of bias and variance/cCRLB ratio in different denoising algorithms. (a, b) Bias and variance/cCRLB ratio plots at different expected photon count levels. Each point and its error bar of the plots were calculated by taking the mean and the standard deviation of bias and variance/cCRLB ratio at different expected photon count levels.

To compare the three denoising algorithms at higher photon count (Supplementary Fig. 1), we increased the peak expected photon counts to be 50 per pixel, with a uniform background of expected photon counts of 10 per pixel. In this setting, the performance of NCS and NLM were similar with their bias and variance increased accordingly with higher photon counts. However, ACsN’s performance improved significantly as the ratio between its variance and cCRLB achieved to 0.97 on average at structure regions and 0.51 on average at the background regions, with relative low absolute bias (~ 0.40 e^-^ on average at structure regions). A possible reason of ACsN’s improved denoising performance on variance is that ACsN not only took the frequency response into consideration, but also recognized the local pattern in the image and used them to denoise similar local patches. However, this assumption wasn’t considered in our cCRLB calculation. At low-photon-counts condition (as shown in Fig. 2), the SNR of the raw images was low such that ACsN may falsely recognized noisy patterns and used them to estimate other patches translating the noise-induced pattern to other image locations which resulted in estimation imprecisions.

Other than photon counting noise (treated as Poisson distribution), sensor noise (e.g. pixel-dependent readout noise of a sCMOS sensor) also affects the performance of denoising algorithm at each pixel. Interestingly, by quantifying the influence of sensor noise on denoising variance lower bound, we found that cCRLB of a single pixel was insensitive towards its intrinsic readout-noise level. This behavior differed from CRLB which increased rapidly with increasing readout-noise standard deviation (SD). To illustrate this observation, we compared cCRLB and CRLB of individual pixels at different locations (Fig. 4). As shown in Fig. 4b, when the pixel has ignorable readout-noise SD (σ=0) and constant gain of 2.17 (Fig. 4b Location 1), the cCRLB was as low as 7.66 (e^-^)^2^, 3.5 times lower than the CRLB (26.57 (e^-^)^2^). When increasing the readout-noise SD of the pixel to 40 ADU, the cCRLB increased to 10.53 (e^-^)^2^ compared to CRLB 491.3 (e^-^)^2^, a 47 times difference. The ratio between CRLB and cCRLB (Fig. 4c) showed that with an increasing readout-noise SD, the benefit of denoising on a single pixel increased. However, such improvement of cCRLB comparing to CRLB in a single pixel came at the cost of slightly increasing cCRLB of its neighbor pixels. Fig. 4c showed that with readout-noise SD of center pixel increased from 0 to 40 ADU, the mean of ratio between CRLB and cCRLB of 8 neighbor pixels decreased from 3.2 to 2.9. Our result showed the possibility of denoising algorithm estimating pixels precisely with high readout noise. This further highlighted the importance of denoising algorithm development in microscopy images, especially for sCMOS sensor where pixel-dependent readout-noise SD varies significantly (e.g. from 1.5 to 40 ADU) among pixels.

**Fig. 4.**
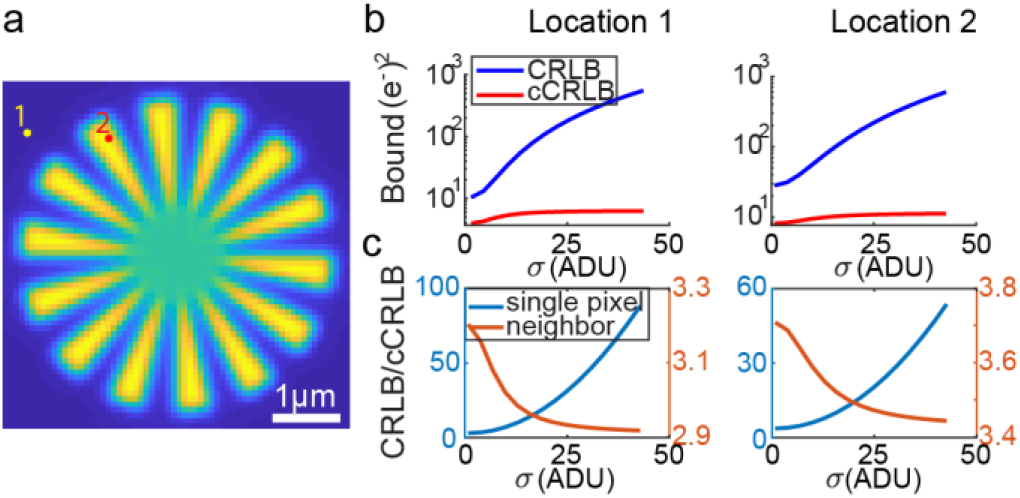
Comparison of CRLB and cCRLB of individual pixels at two locations. (a) The ideal image with the marked locations. (b) Comparisons between CRLB and cCRLB with respect to increasing readout-noise SD in the marked locations. (c) The ratios of CRLB/cCRLB of the center pixel and the mean ratios of its neighbors (8 surrounding pixels).

Further, we demonstrate that cCRLB was highly related to the numerical aperture (*NA*) as well as the effective camera pixel size in the specimen. We expressed the radius of the OTF boundary in the unit of numbers of pixels in the Fourier space as

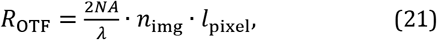

where *n*_img_ is the number of pixels of the image in one lateral dimension (assuming a square image), and *I*_pixel_ is the effective camera pixel size (the physical length of one pixel occupied in the specimen). Therefore, increasing *NA* increases the radius of the OTF boundary causing fewer number of constraints imposed on parameters. As cCRLB depends on the number of constraints and the number of pixels within the image, larger *NA* results in larger cCRLB (Fig. 5). An extreme case will be when the *NA* is sufficiently large such that OTF occupies the entire field of the Fourier transform of the image. In such a case, there is no constraint imposed on the parameters and cCRLB will be equal to CRLB.

**Fig. 5.**
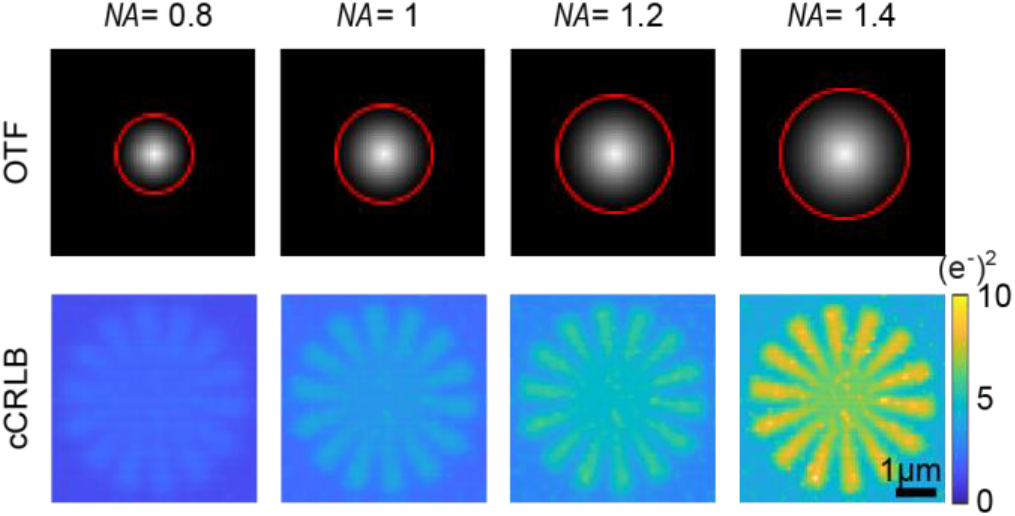
The relationship between *NA* and cCRLB. The red circle on the OTF indicates the OTF boundary.

Next, we investigated the influence of effective pixel size on cCRLB. In order to maintain the size of the field of view as well as the photon flux emitted per area, we changed pixel size while adjusting number of pixels of the image and the expected photon counts per pixel accordingly. Here, we calculated cCRLB in two settings: one with an image of 128*128 pixels, a pixel size of 40 nm, peak photon counts of 5 per pixel, and background photon counts of 2.5 per pixel and the other with an image of 64*64 pixels, a pixel size of 80 nm, peak photon counts of 20 per pixel, and background photon counts of 10 per pixel. In order to compare these two cCRLB maps, we binned the cCRLB map of 128×128 pixels into a 64×64 pixels map (Fig. 6a). The binned cCRLB represents an approximation of the variance lower bound of binned denoised image. As shown in Fig. 6, we found that the cCRLB with the pixel size of 40 nm are four times smaller than that of 80 nm on average. This observation suggested that choosing a smaller effective pixel size during microscopy experiments benefited denoising performance with a lower achievable variance bound.

**Fig. 6.**
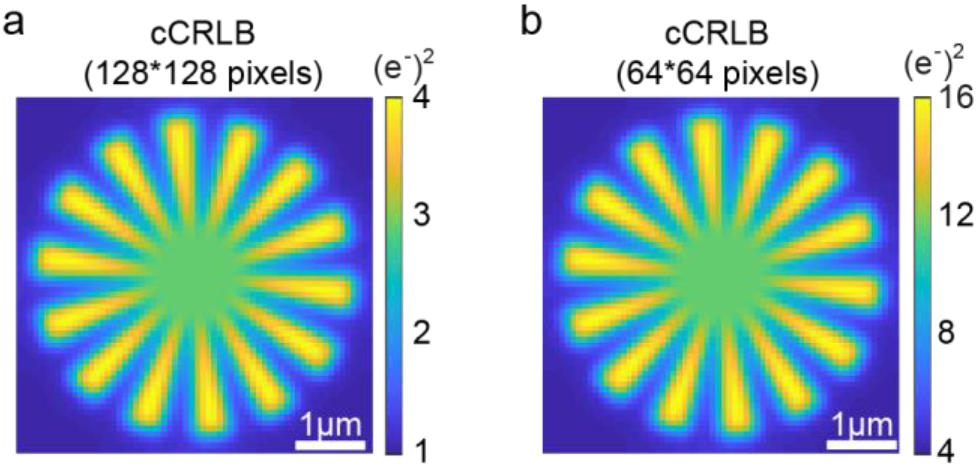
Comparison of cCRLB maps calculated from different pixel sizes while photo flux per area remains constant. (a) cCRLB map was calculated from a simulated image of 128×128 pixels with the pixel size of 40 nm and binned to 64×64 pixels. (b) cCRLB map was calculated from a simulated image of 64×64 pixels with the pixel size of 80 nm. In these two maps, the underlying structure, the field of view, and photon count per area were kept the same. Here, *NA* was set to 1.4, the wavelength of light was set to 700 nm and a uniform readout-noise SD was assumed across the field of view at 3.46 ADU with a uniform gain of 2.0.

cCRLB represents the theoretical lower bound on variance of unbiased estimation of microscopy images. However, to approach unbiased estimation, most denoising algorithms will have to sacrifice their ability of reducing estimation variance (Fig. 7). For example, NCS algorithm has an inherent parameter *α* serving as a balancing parameter between its likelihood function based on the noise model and the prior knowledge. By tuning this parameter, one can balance the tradeoff between bias and variance. We demonstrated this effect in Fig. 7. When *α* = 0.1, we observed the denoised image with relative small bias while its variance performance was far from the theoretical bound (cCRLB). However, as parameter *α* increased to 3, the variance of denoised image had reached to 7.04 (e^-^)^2^ compared to variance of original noisy image 14.89 (e^-^)^2^ while introducing the bias with ~0.28 e^-^ on average at structure regions and the ratio between variance and cCRLB was 1.54 on average. As this parameter further increased (*α* = 10 and *α* = 100), the variance was reduced to 3.55 (e^-^)^2^ and 1.70 (e^-^)^2^ on average respectively, at the same time, the absolute bias at the structure regions increased to an average amplitude of 0.35 e^-^ and 0.5 e^-^.

**Fig. 7.**
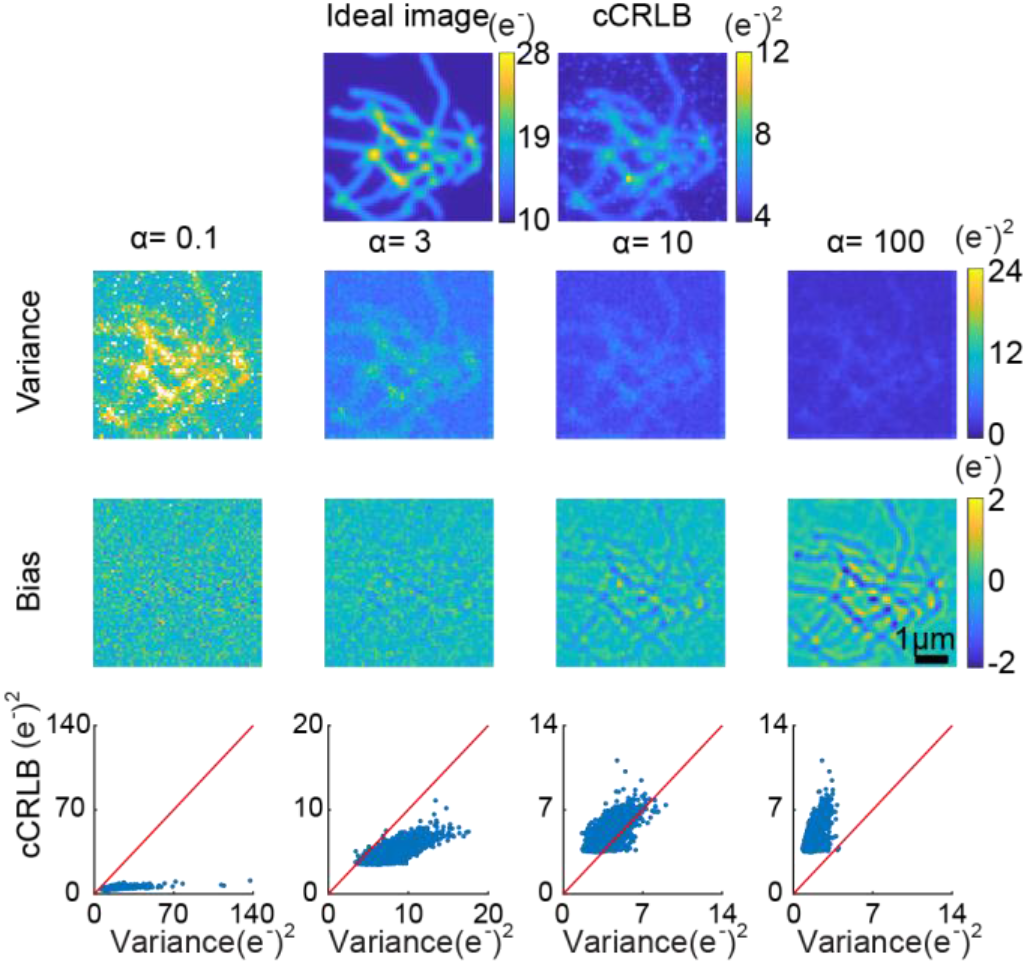
Trade-off between variance and bias of NCS denoising algorithm. Here we applied a simulated microtubule image for denoising demonstration. The variance maps (second row) and bias maps (third row) were calculated by using different parameter *α* in NCS. Every pixel’s cCRLB and its denoising variance were shown in the scatter plot (fourth row) with the solid red line representing the cases where cCRLB and the estimation variances are equal.

Our development of cCRLB provides a theoretical lower bound on the estimation variance of unbiased denoising algorithms for microscopy images by considering the finite spatial frequency response in a microscope system. This work provides a general framework to incorporate additional knowledge, either from the imaging system or potentially from the specimen, to estimate the precision performance limit of the denoising estimation by formulating a set of constraint equations representing the prior knowledge. In addition, by using cCRLB, we show that the readout statistics (readout-noise SD) of individual pixel has little influence on how precise the pixel intensity can be estimated, suggesting the importance of developing denoising algorithms especially for camera sensors with readout-noise variation in different pixels (e.g. CMOS sensor). Although cCRLB does not guarantee the existence of an estimator or an algorithm achieving such limit, we expect cCRLB will be a useful tool predicting the variance limit for a proposed microscopy denoising algorithm.

## Acknowledgment

We would like to thank Fan Xu for the suggestions on our simulations, fruitful discussions and helps on revising the manuscript. We would also like to thank Benjamin Brenner for the suggestions on our simulations and manuscript.

