## Supplementary Fig. 1 for "Variance lower bound on fluorescence microscopy image denoising"

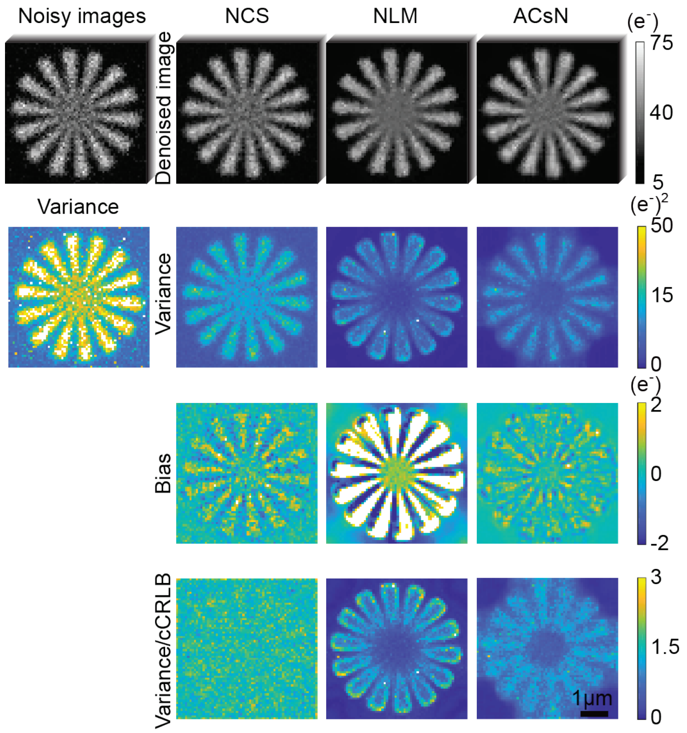


Supplementary Fig. 1. Comparison of three different denoising algorithms at high photon count case. First row shows examples of noisy image (raw image before denoising) and denoised images from NCS, NLM and ACsN algorithms. Variance map (second row) and bias map (third row) were calculated using 100 frames of corresponding noisy or denoised images. Variance/cCRLB ratio maps (fourth row) were calculated from variance maps divided by cCRLB pixel by pixel.
